# Unravelling the origins of boldness: A common garden experiment with cave fish (*Barbatula barbatula*)

**DOI:** 10.1101/2023.10.05.561011

**Authors:** Jolle W. Jolles, Alexander Böhm, Alexander Brinker, Jasminca Behrmann-Godel

## Abstract

Many animals show an aversion to bright, open spaces, with significant variability seen across species, populations, and individuals within populations. Although there is much interest in the underlying causes of this behaviour, few studies have been able to systematically isolate the role of heritable and environmental effects. Here we addressed this gap using a common garden experiment with cave fish. Specifically, we bred and cross-bred cave loaches (*Barbatula barbatula*) in the lab, raised the offspring in complete darkness or normal light conditions, and studied their light avoidance behaviour. Cave fish spent much more time in a light area and ventured further out, while surface fish spent considerable time on the edge between light and dark areas. Hybrids behaved most similarly to cave fish. Light treatment and eye size and quality only had a modest effect. Our results suggest light avoidance behaviour of cave fish has a heritable basis and is fundamentally linked to increased boldness rather than reduced vision, which is likely adaptive given the complete lack of macropredators in the cave environment. Our study provides novel experimental insights into the behavioural divergence of cave fish and contributes to our broader understanding of the evolution of boldness and behavioural adaptation.

## 1. Introduction

Throughout their life, animals must continuously try to acquire resources such as food and mates while minimizing potential risks, including predators, aggressive conspecifics, and detrimental environmental conditions. This dynamic interplay of risk-taking and avoidance, also known as boldness [1], needs to be flexible to some degree for animals to optimally fine-tune their behaviour in response to changes in their environment. But in the long run, when animals of different species, or local populations within species, experience consistently different environmental conditions that influence this trade-off, differences in average reaction norms are expected to evolve.

Predation risk is one of the major environmental conditions thought to shape boldness behaviour. Compelling evidence comes from manipulative selection experiments, with bold lizards being favoured on predator-free islands and shy individuals the most likely to survive in the face of predators [2]. Also, numerous studies exist that report differences in average boldness between populations that differ in predation pressure, with animals from low predation sites tending to be bolder [3–6]. But while this is suggestive of heritable differences in boldness [7], it is often challenging to properly isolate such effects from behavioural plasticity and experience [8]. This can be overcome by common-garden experiments, whereby animals from different populations are reared under identical laboratory conditions [9]. However, as populations may differ in numerous other uncontrolled variables, such as animals forming part of complex multi-species communities, and hybridization may occur, it often remains difficult to pinpoint the exact drivers of selection.

Underground cave systems, generally characterised by perpetual darkness, limited primary production, and scarcity of food, exist across the world [10,11]. Despite these challenging conditions, subterranean habitats host a broad array of life, including over 200 cave-dwelling fish species [12–14] that show remarkable phenotypic convergence, including reduced eyes and pigment [10,11] and a spectrum of behavioural adaptations [8,15–21]. In contrast to surface (epigean) environments, the subterranean realm provides a highly constrained range of environmental conditions marked by high stability, predictability, and homogeneity, thereby providing a clear stage for selection [11]. Moreover, the presence of closely related surface populations enables direct comparisons. Hence, cave fish represent a powerful model for investigating the evolution of behavioural traits in response to environmental change [22,23] and analysing adaptive changes driven by well-defined selection pressures [24].

Already considerable work has investigated the behavioural adaptations of cave fish, including in terms of foraging behaviour, schooling, mate choice, aggression, and movement kinematics [8,15–21]. Although a number of studies have documented cave fish to have a reduced stress response [25] and show less light avoidance [26–28], the shy-bold axis [29] has so far received little focused attention in cave fish. This is clearly reflected by its lacking in a recent primer on cave fish and a review on their behaviour [24,30]. This is surprising given that the cave environment tends to completely lack macropredators, providing a potential strong selection pressure for increased boldness behaviour for cave fish.

The aim of this study was to systematically investigate the evolutionary drivers of boldness using cave fish as our model system. We thereby take advantage of a recently discovered cave population of stone loach (*Barbatula barbatula*), the world’s northernmost known cave fish [31]. These fish inhabit the Danube-Aach system, an underground karst water system in Southern Germany, and exhibit typical morphological traits for cave organisms, such as diminished eyes and pale body colouration (see Figure 1A). Despite likely only having evolved in the last 20.000 years, the cave population is genetically isolated from surface populations, as revealed by microsatellite analyses [31].

**Figure 1.**
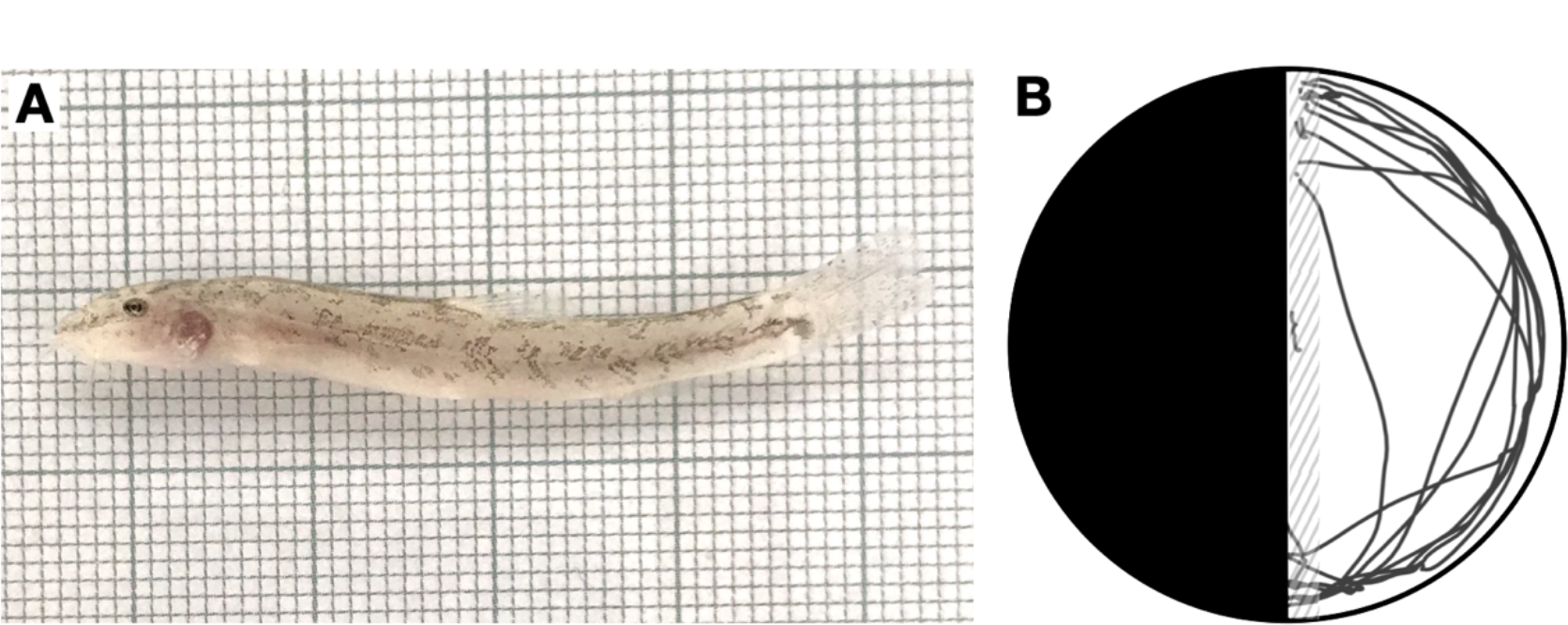
A) Photo of an experimental fish with cave origins from the DL treatment. B) Schematic drawing of the experimental arena with the tracking data of one randomly selected trial overlaid, with the dashed area indicating the area in which fish were labelled as ‘partly out’ and beyond as ‘fully out’.

To determine the role of genetic *versus* environmental effects underlying light-avoidance behaviour, we successfully established a laboratory population of cave loaches and conducted a common-garden experiment. We thereby cross-bred cave and surface fish and raised the resulting offspring – cave, surface, and hybrid fish – in either complete darkness or under normal day-night conditions. We then subjected the fish to a light-dark preference test, a well-established assay that consists of one light and one dark half, and measures fishes’ tendency to avoid the light area, which is naturally aversive for many animals, including fish [32]. By furthermore conducting detailed behavioural tracking and morphological analyses, we were able to disentangle the potential proximate mechanisms unerlying the observed behavioural differences. By using cave loach as our model system, our study helps better understand the behavioural differentiation of cave fish and provides new insights into the evolution of boldness behaviour.

## 2. Material and methods

### 2.1 Study Animals

We captured stone loaches from the Danube-Aach cave system in southern Germany with the help of a professional cave diver, as well as from two nearby surface populations by electrofishing (described in detail in [31]). Fish were moved to our lab at the Limnological Institute of Konstanz and housed in 60 L tanks, containing a substrate of sand and gravel, stones, a PVC pipe for shelter, and an air stone for aeration. Cave and surface fish were kept in separate rooms with similar conditions, except for the light treatment, with cave fish kept in complete darkness and surface fish under normal light-dark conditions. Tanks received constant flow-through of filtered water from Lake Konstanz, and fish were fed *ad libitum* a combination of dry food, frozen bloodworms, and live *Daphnia*.

### 2.2 Common garden experiment

After more than six months of acclimation in the lab, we conducted a controlled pairwise breeding experiment whereby we paired fish with cave, surface, and mixed origins to establish a laboratory-reared population of fish with different origins (see [33] for further details). Although breeding was challenging, we were able to obtain a large number of eggs from the different pairs. Upon detection, we immediately separated them into two batches, one that we kept in complete darkness (DD), simulating cave conditions, and one under natural light-dark conditions (DL). The eggs were let to hatch in 20 L circular tanks with water on a flow-through system. After hatching, the fry – from now on termed ‘cave fish’, ‘surface fish’, and ‘hybrids’ based on the origins of their parents – were fed fresh artemia and crushed flake food (Tetra) *ad libitum*. As fish grew, food particle size was increased and trout pellet food (INICIO plus trout, 0.3 mm, BioMar), live Daphnia, and frozen bloodworms were provided.

### 2.3 Behavioural experiment

When the fish were about two months old, we netted them individually from their holding tanks, anaesthetized them with MS-222 (0.1 g/L), and moved them to our experimental room. Immediately, each fish was photographed from above in a Petri dish to measure body length before moving them to an individual holding tank, where they were allowed to wake up from anaesthesia. Tanks were on a flow-through system, with fish kept in individual compartments (20 cm × 10 cm; 10 cm water depth), each with an artificial plant for shelter, separated by transparent, perforated walls to enable the transfer of visual and chemical cues. The experimental room had fluorescent lighting on a normal day-night cycle.

After a day of acclimation, we subjected the fish to the experimental arena, a circular glass tank (30 cm diameter, 10 cm water depth; Fig. 1B) with one dark and one light half, also sometimes referred to as the half-moon assay [34], made with black and white laminated paper covering the bottom and sides of the tank. Fish were thereby gently taken from their individual holding compartment using a dipnet and released into a transparent cylinder in the centre of the arena where they were left to acclimate. After two minutes, the cylinder was raised and the trial began. At the end of five minutes, the trial ended and the fish was gently moved back to its holding compartment.

In total, we used 8 testing arenas to facilitate a high throughput of animals. The arenas were positioned in a large experimental area that provided diffuse lighting from above and was surrounded by opaque curtains to minimize external disturbances. Between trials, arenas were rotated up to 45 degrees to avoid any possible localization effects. Each fish was tested twice, once in the morning and once in the afternoon using the same testing order but a different randomly selected experimental arena. Trials were recorded at 24 frames per second using four Raspberry Pi computers and associated HD cameras (one camera per two tanks) on a local area network [35] and *Pirecorder* software was used to facilitate controlled and automatic video recording and file organisation [36]. After the behavioural experiment, fish were euthanized with an overdose of MS-222 (3g/L), transferred to formalin (4%), and photographed laterally and dorsally using a binocular microscope for detailed morphological measurements.

### 2.4 Data processing

Videos of the behavioural trials were automatically tracked using custom tracking software in Python (ATracker). Due to the dark background of the black half of the testing arena, we were only able to accurately track the fish in the white half of the arena. We thereby used background subtraction methods and blob detection, tracking the centroid of the fish and using an object area roughly corresponding to half that of a fish as the minimum threshold size for tracking. After tracking, the data for each trial was carefully checked by observing trajectory-overlaid videos. If needed, a trial was either re-tracked with adjusted parameters or the erroneous sections fixed by manual tracking using a custom user interface. Subsequently, all tracking data was processed in R [37] using RStudio. We thereby computed, for each trial, the proportion of time a fish was ‘out’ in the light area, was out but still near (object centroid within 15 mm of) the dark area (‘partly out’, see Fig. 1B), and tracked but ‘fully out’ (i.e. further than 15 mm from the dark area). Finally, we computed each fish’s median speed when fully out, allometrically corrected (i.e. in body lengths per second; BL/s). Morphological measurements were acquired from the lateral photos of the formalin-fixed fish using ImageJ [38]. Specifically, the eye and lens diameter to the nearest μm. We furthermore classified lenses into four classes to indicate their quality: Class 3 (normal), lens uniform in color with a diameter approximately half that of the eye; Class 2, lens cloudy or milky, diameter 80-100% of Class 3 lenses; Class 1, lens < 80 % the size of Class 3, completely white; and Class 0, no lens detected. Diversification of the complete morphology of cave loaches was the focus of another study and is described in detail there [33].

### 2.5 Data analysis

We used a linear mixed modelling approach using backward stepwise elimination to investigate the extent that behaviour in the light-dark preference test was correlated with fish’s origin (cave, surface, hybrid), raising treatment (DD, DL), and eye morphology (lens quality and relative lens diameter). Fish ID, camera (1-4), and trial (1-2) were added as random factors. We visually inspected residuals to ensure homogeneity of variance, normality of error, and linearity, and square-root transformed data where necessary. Post-hoc pair-wise comparisons were done using the Tukey method to adjust p-values for multiple comparisons using the *emmeans* package v1.8.3 [39]. Calculations of repeatability were performed with the *rptR* package in R [40]. All data were analysed in R 4.2.2 [37]. In total, we tested 192 fish, of which 50 were cave fish, 42 hybrids, and 100 surface fish, with each fish tested twice. However, 15 trials had to be excluded due to issues with video recording or incomplete morphological measurements.

## 3. Results

On average, fish showed a tendency to avoid the light area and spent on average 22% ± 1% of their time out in the white half of the arena. This behaviour was highly repeatable across the two trials (*R* = 0.60 [95% confidence intervals: 0.51-0.69]), also when considering fish of cave origins only (*R* = 0.57 [0.33-0.73]). The time fish spent in the light area was strongly linked to fish’s origin (χ^2^ = 92.30, *p* < 0.001; Figure 2A). Cave fish spent significantly more time in the light area than surface fish (*t*_183_ = 10.09, *p* < 0.001). Hybrids resembled this behaviour, with only a trend for cave fish to go out more (*t*_183_ = 2.22, *p* = 0.071) and hybrids spending significantly more time out than surface fish (*t*_185_ = 6.93, *p* < 0.001). The effect of origin was even stronger when considering the time when fish were fully out (i.e. further away from the dark area; χ^2^ = 99.77, *p* < 0.001). Also raising treatment had a significant effect on fish’s time spent in the light area, with those fish raised in complete darkness spending more time out than fish raised under normal light-dark conditions (χ^2^ = 10.32, *p* = 0.001; Figure 2A). However, its effect size was limited, at only 10% of the effect of fish’s origin.

**Figure 2.**
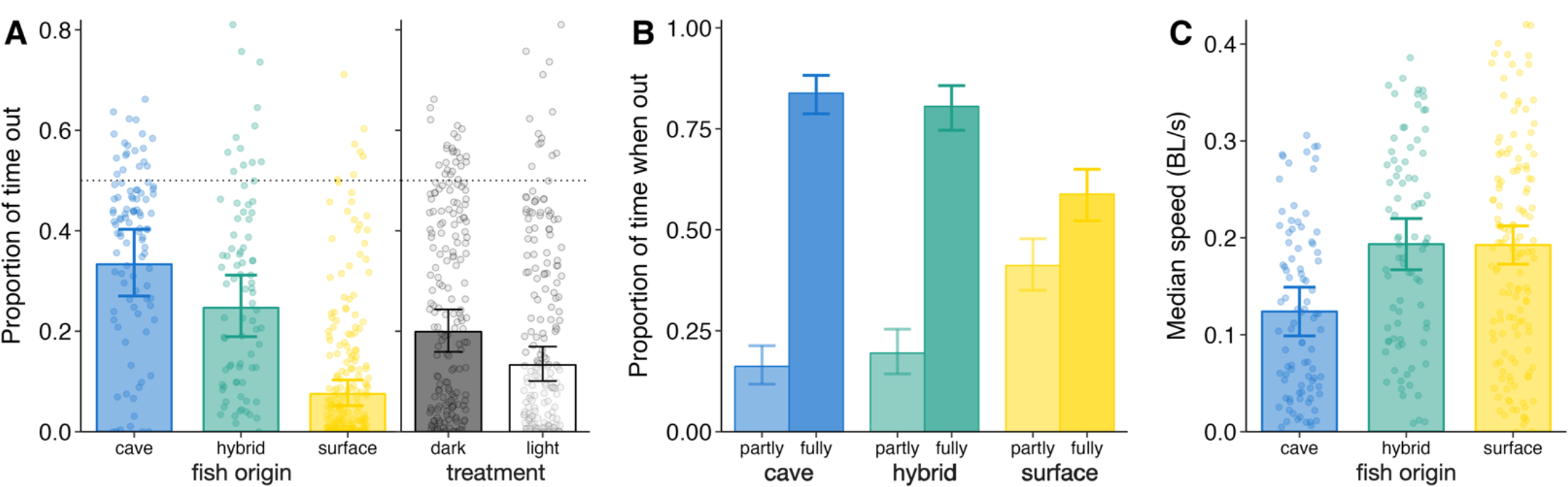
A) Bar plot of the proportion of time fish were out in the light area for the different origins and two raising treatments. B) Barplot of the relative proportion of time that fish were either partly, i.e., tracked within 15 mm from the ark area, or fully out for the different origins. C) Fish’s median speed when fully out for the different origins. Bars and errors are based on model output, back-transformed where applicable, and points show the raw data.

Of the time fish were tracked in the light area, they spent considerable time lingering on the edge with the dark area, on average 33.7 % ± 1.6 %, often with much of their body still concealed. There was large variability in this behaviour, which was strongly positively linked to the fish’s origins (χ^2^ = 61.35, *p* < 0.001; Figure 2B): surface fish spent almost half of their time out while being partly concealed, which was more than double that observed for cave and hybrid fish (*t*_172_ = -7.77, *p* < 0.001; *t*_174_ = -6.15, *p* < 0.001 respectively). As this effect could possibly be influenced by the proportion of time fish were out in the white half, we re-ran the analysis with a subset of the data where fish spent at least 15s out. This did not qualitatively change the effect (χ^2^ = 32.06, *p* < 0.001, *n* = 263). Also raising treatment had no significant effect on the time fish spent partly out in the light area (χ^2^ = 0.52, *p* = 0.472).

Fish’s median movement speed (when fully out) was also linked to fish’s origin (χ^2^ = 19.93, *p* < 0.001, Figure 2C): on average, cave fish swam significantly slower than surface fish (t_163_ = -4.23, *p* < 0.001) and hybrids (t_152_ = -3.75, *p* < 0.001). There was no difference in swimming speed between surface fish and hybrids (*t*_163_ = 0.05, p = 0.998). Raising treatment again had no significant effect (χ^2^ = 1.65, *p* = 0.199).

In terms of eye morphology, there was large variability in lens quality among the experimental fish: all fish with surface origins had perfect eyes (class 3), and so did 95% of fish with hybrid origins, but cave fish varied considerably in lens quality with only 2% having perfect eyes, 36% having lower class 2 eyes, 50% having class 1 eyes, and 12% having no lens at all. Overall, lens quality was a good predictor of fish’s time spent in the white half of the arena (χ^2^ = 52.17, *p* < 0.001; Figure 3A), with fish that had lower-quality lenses spending more time out. However, this effect was strongly linked to the variation in lens quality linked to origin, and was not significant when considering cave fish only (χ^2^ = 5.73, *p* = 0.125). Similarly, considering relative lens diameter instead, there was an overall effect that cave fish with smaller lenses spent more time out in the light area (χ^2^ = 4.83, *p* = 0.028). But this effect was strongly influenced by fish without any eye lenses behaving at chance level (*t*_11_ = 0.66, *p* = 0.520; see Figure 3B), and excluding those fish resulted in a non-significant effect (χ^2^ = 0.14, *p* = 0.707). Relative lens diameter was not correlated with the observed variation in movement speed among the fish (χ^2^ = 0.02, *p* = 0.884).

**Figure 3.**
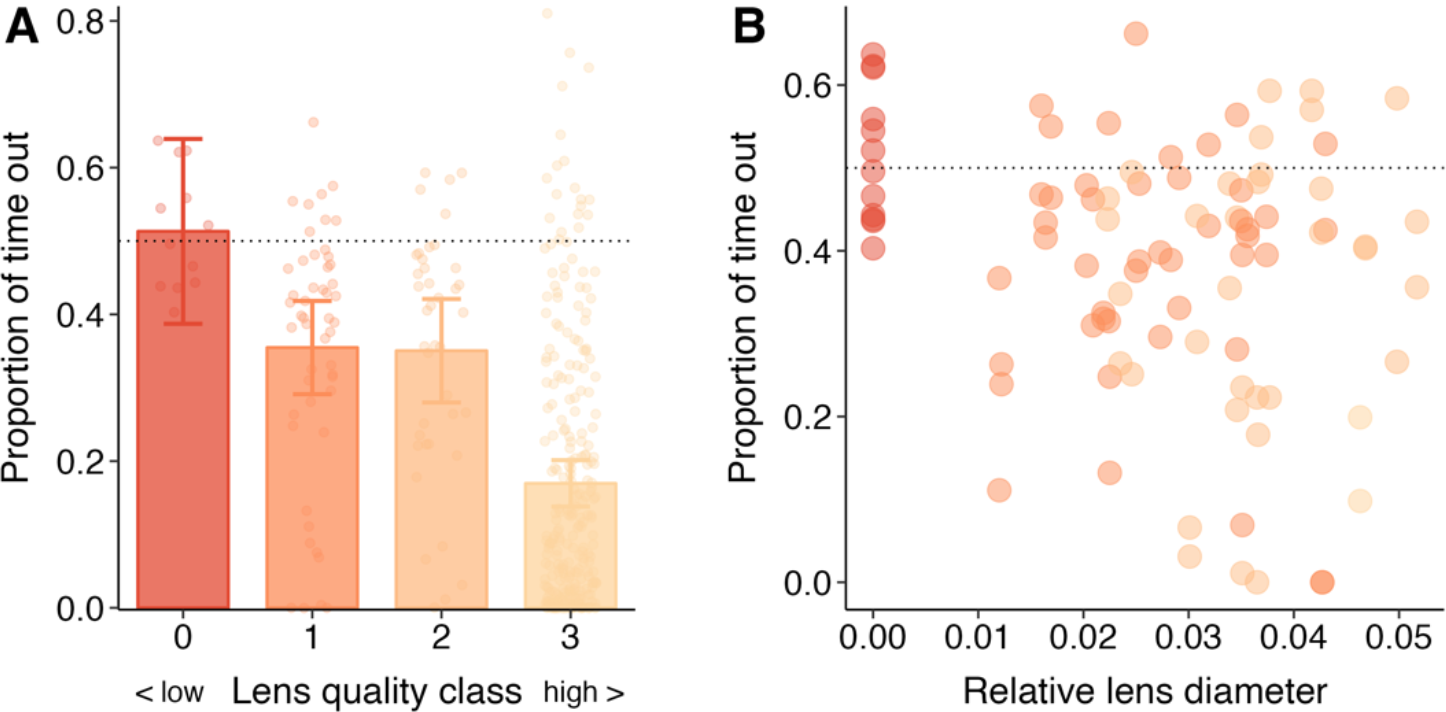
A) Bar plots of the proportion of time that fish with cave origins were out in the white half of the arena categorised based on lens quality (class 0-3 with 0 indicating no lens and 3 an eye with perfect lens). Bars and errors are based on model output; points show the raw data. B) Scatterplot of fish’s proportion of time in the white half as a function of the diameter of their eye lens relative to the length of their head (cave fish only). Point colour indicates eye quality class (as in A). Dotted lines indicate chance level, i.e. fish would spend 50%/50% of their time in the black/white half.

## 4. Discussion

In this study we conducted a comprehensive investigation into the factors influencing light avoidance behaviour in European cave loach, combining a common garden experiment and detailed behavioural and morphological analyses. We found that laboratory-reared fish with cave origins spent significantly more time in a bright open area than offspring with surface origins. This outcome is consistent with previous research highlighting reduced light avoidance behaviour by cave fish. Rearing fish in either complete darkness or under normal day-night conditions within a common garden setting revealed light exposure only explained only a small proportion of the behavioural variance observed. Furthermore, detailed tracking of the fish revealed another clear difference between the fish: surface fish spent a considerable proportion of their time lingering between the light and dark areas, something not seen for cave fish. Overall, the behaviour of hybrid fish most closely resembled that of cave fish, emphasizing a heritable origin for the behaviours observed. To investigate the potential of reduced vision as a proximate driver for the observed behaviour, we looked at the size and quality of the fish’s eye lenses. This showed that while fish completely lacking eye lenses spent equal amounts of time in the light and dark areas, for fish containing eye lenses (the far majority), the size or quality of the eye lens did not correlate with time spent in the light area. Together, our results strongly indicate that the light avoidance behaviour of cave fish is closely linked to boldness and largely has a genetic origin. These findings not only contribute valuable insights into the behavioural divergence of cave fish, they have broader implications for our understanding of behavioural adaptation and the origins of boldness behaviour.

The most prominent behavioural difference we observed among the laboratory-reared fish was the time they explored the white half of the arena, with offspring with pure surface origins showing three times more avoidance of the light area than fish with pure cave origins. Our finding is in line with stone loach naturally avoiding brightly lit areas [41] and previous work showing a generally reduced light avoidance tendency in cave fish [27]. Notably, however, there is no clear consensus, with some studies reporting no difference with surface fish [25,27] and others describing strong photophobic responses in cave fish [28] – even for fish lacking eyes completely [42]. For an animal that lives in complete darkness, responsiveness to light may not confer any advantages and may be hypothesized to erode away over time. However, rather than responsiveness to light itself, the observed difference in light avoidance behaviour may reflect an underlying difference in risk avoidance behaviour as animals tend to prefer dark places to hide from predators. In other words, the tendency of cave fish to venture out and explore a light area may reflect boldness behaviour [43]. This is supported by the finding that pharmacological agents can modulate light avoidance behaviour [25,31] and correlates with other classic boldness tests [44,45]. Further support for cave fish to be bolder comes from our detailed tracking: cave fish swam slower and spent more time further away from the dark area, while surface fish spent a considerable part of their time lingering on the edge between the black and white half, often with most of their body still being concealed.

This behaviour may not only reflect higher risk avoidance of surface fish, because risk is generally perceived to increase with distance from a refuge [46], but also increased risk-assessment behaviour, which animals tend to do on the border between the safety of a refuge and a more risky area [47]. Intriguingly, it is possible that the reduced schooling behaviour of cave fish [15,27,48] is also a reflection of increased boldness as it tends to be negatively linked with attraction to others [49] and grouping tends to increase with predation pressure [50]. Although ‘boldness’ is the most studied animal personality trait [29,51] and we found light avoidance behaviour to be strongly repeatable across the two trials, this is not the focus of our study but our intention to better understand the drivers of boldness in the general sense of increased risk-taking behaviour [1].

In contrast to other systems, for cave fish there is a potential alternative explanation than boldness for reduced light avoidance behaviour: that of impaired vision [27,34]. Eye regression is one of the most prominent traits of cave fish [10,11,30], and hence cave fish may simply not be able to see their environment properly. For the population of cave loaches studied, the adaptive process has not yet resulted in complete eye loss. For the few fish that did not have any eye lenses (about 1/10th of the offspring with cave origins), we did observe they spent equal amounts of time in the dark and light areas, in line with previous work [25,27]. However, for the rest of the fish with cave origins, the relative size of the lens did not correlate with the time spent in the light area (but see [27]). This suggests that, although a complete lack of vision may help explain a lack of light avoidance, variation in reduced vision explains little of the variation in light-avoidance behaviour overall (see also [26]) and is not likely to not be the primary underlying mechanism. The most compelling evidence comes from the behaviour of the hybrid fish, as despite almost all having normal eyes lenses, there was a large difference between hybrids and surface fish, with hybrids spending significantly more time out in the light area.

As part of our common garden design aimed at disentangling genetic *versus* environmental effects, we raised fish in either complete darkness or a natural day-night schedule and otherwise identical conditions. Although raising treatment had a significant effect on light avoidance behaviour, with fish raised in complete darkness spending more time in the light area, the effect was relatively weak, at 10% of that of origin. Together with the finding that hybrid fish behaved most like the offspring with pure cave origins, these results suggest that the observed difference in boldness behaviour between the cave and surface fish largely has a genetic origin and might be largely fixed in the cave fish population. As the cave environment provides a constrained range of highly stable environmental conditions [11], there are only a few candidate variables that may have led to this divergence, in particular that of predation risk. Namely, underground caves tend to lack macropredators completely, which would ease selection for risk-avoidance behaviours such as hiding: the increased boldness of cave-dwelling fish may be adaptive as being more exploratory in a completely dark environment can help increase the likelihood of encountering food or mates. If reduced predation risk is indeed the evolutionary driver for increased boldness in cave fish, one would also expect anti-predator behaviours to be reduced. There is indeed some support for this, including reduced startle responses in cave fish [52] and generally lower responsiveness to stressful stimuli [15,25,27]. Although Bierbach et al. [53] reports similar levels of anti-predator behaviour in cave- and surface populations of Mexican Mollies, they argue this is likely related to some level of predation still being present. More work is needed to investigate the generality of increased boldness and anti-predator behaviour in cave fish. Another potential driver of increased boldness in cave fish is low resource availability [15]. However, we believe it not to be the primary factor, at least in cave loach, as the large catchment (250km^2^) and major inputs of water via Danube sinks [31] are expected to ensure a relatively good food supply. This is indeed reflected by the frequent observation of planktonic copepods and small crustaceans in the stomach contents of freshly caught cave fish (own observations), with the latter also frequently observed in the caves (pers. comm. of the cave divers). We cannot exclude that the increased boldness behaviour of cave fish is the result of pleiotropic effects or drift of random mutations not counter-selected for in the cave environment [54]. But given the adaptiveness of boldness behaviour in the context of extended exploration for food and mates, we expect the influence of such effects to be rather small.

While cave fish across the world vary widely in their degree of phenotypic change [10], only a couple of species have been extensively studied (in particular *A. mexicanus* [13,30]). Studies like ours on a much less studied (and Europe’s only known) species are therefore needed to validate the existence of universal patterns, or reasons for lack thereof, across cave-dwelling fish. Furthermore, by focusing on a relatively recently diverged cave fish population, our study may help better understand the circumstances and evolutionary timescales over which such phenotypic changes may occur. Our study also provides new insights into behavioural evolution more generally. Namely, while a number of studies exist that document fish from sites with high predation pressure to be bolder than those with low or no predation pressure [3,5,6,55], often genetic and environmental effects cannot be properly disentangled. While common garden experiments can provide a solution, a multitude of variables may differ between study populations (see e.g. [4]), making it hard to pinpoint the selection pressures that led to phenotypic change. Fish populations in caves live in a highly stable and low complexity environment that completely lacks any macropredators. Therefore, our comparison of the behaviour of cave and surface fish provides new evidence that predator risk is a strong selection pressure underlying the evolution of boldness behaviour.

In conclusion, our study, which combines a common garden experiment with movement tracking of European cave loach in a classic light-dark preference test, sheds light on the remarkable behavioural evolution of cave fish. Despite its relatively recent origin, dating back as little as 20.000 years ago [31], we demonstrate how European cave loach have diverged considerably in terms of light avoidance behaviour, and show this behaviour has a heritable basis and is fundamentally linked to boldness. Beyond its implications for our understanding of cave fish evolution, our research offers broader insights into the origins of boldness and behavioural adaptation. Our study shows that cave fish provide a valuable model for studying patterns of behavioural evolution, and hope our work inspires future studies to unravel its underlying complexities.

## Ethics

This study was carried out in accordance with the Protection of Animals Act Germany. All of the animals were maintained and handled according to the Institutional Animal Care guidelines of the University of Konstanz. Animal collection was approved by the regional council of Tübingen (reference number: 33-4/9220.51-3) and animal holding and experimentation was approved by the regional council of Freiburg (reference numbers: 35-9185.64/1.1 and 35-9185.81/G-19/30).

## Data accessibility

Data accompanying this paper can be found on the Open Science Framework: https://osf.io/mgeku/.

## Authors’ contributions

JB-G conceptualized the overall study and oversaw the breeding of the fish; JWJ, ABö. and JB-G designed the behavioural experiments; ABö ran the experiments supervised by JB-G and JWJ; JWJ conducted the tracking of the behavioural trials; ABö and JB-G handled and photographed the fish and conducted morphological analysis together with ABr; JWJ analysed the data and wrote the manuscript. All authors commented on subsequent versions and gave final approval for publication.

## Acknowledgements

We thank cave diver Joachim Kreiselmaier and his team for their invaluable help with catching the cave loach under extremely challenging conditions and Myriam Schmidt for her help with fish maintenance.

